# Phenotypically wild barley shows evidence of introgression from cultivated barley

**DOI:** 10.1101/2024.07.01.601622

**Authors:** Chaochih Liu, Li Lei, Mingqin Shao, Jerome D. Franckowiak, Jacob B. Pacheco, Jeness C. Scott, Ryan T. Gavin, Joy K. Roy, Ahmad H. Sallam, Brian J. Steffenson, Peter L. Morrell

## Abstract

Plant conservation hinges on preserving biodiversity, which is crucial for long-term adaptation. Multiple studies have reported genetic evidence of crop-to-wild introgression in phenotypically wild accessions of wild barley (*Hordeum vulgare* ssp. *spontaneum*). We examined 318 Wild Barley Diversity Collection (WBDC) accessions for evidence of introgression from cultivated barley. Using SNP genotype and exome capture data, we performed local ancestry inference between the 318 WBDC accessions and cultivated barley to identify genomic regions with evidence of introgression. Using the genomic intervals for well-characterized genes involved in domestication and improvement, we examined the evidence for introgression at genomic regions potentially important for maintaining a wild phenotype. Our analysis revealed that nearly 16% (48 of 318) of WBDC accessions showed evidence of introgression from cultivated barley, and up to 16.6% of the genome has been introgressed. All accessions identified as introgressed based on domestication-related phenotypes show clear genetic evidence of introgression. The size of runs of identity by state and local ancestry inference suggests that most introgression did not occur recently. This study suggests a long history of genetic exchange between wild and cultivated barley, highlighting the potential for introgression to influence the genetic makeup and future adaptation of wild populations, with implications for plant conservation strategies.

## Introduction

*Ex situ* conservation is essential for preserving diversity in many animal and plant species in the long-term. Plant conservation efforts have focused on maintaining collections of important crops and their wild relatives. These resources are essential to meet the changing demands of crop productivity, sustainability, and resilience of agriculture to added challenges such as climate change (Schoen & Brown 2001; Bohra *et al*. 2022; Dempewolf *et al*. 2023; Flint-Garcia *et al*. 2023; McCouch & Rieseberg 2023). Preservation of natural habitats for the *in situ* conservation of crop wild relatives offers many advantages, but individual natural populations can be readily lost to land development, disturbance, or changing climate. *Ex situ* conservation offers the further advantage of permitting phenotypic characterization in a common garden environment and the accumulation of organized information collected about accessions (e.g., diversity levels, traits, characteristics, etc.). This information is particularly valuable for researchers and plant breeders to make informed decisions when selecting material for use (Dempewolf *et al*. 2023). Introgression between crops and their wild relatives is common (Rieseberg & Wendel 1993; Ellstrand 2003; Ellstrand *et al*. 2013) and sometimes adaptive (Morrell *et al*. 2005; Hufford *et al*. 2013).

Wild barley (*Hordeum vulgare* ssp. *spontaneum*) and cultivated barley (*Hordeum vulgare* ssp. *vulgare*) are most readily distinguished by the lack of shattering of rachis in the cultivated form; they are cytogenetically indistinguishable, cross readily and commonly hybridize in natural settings (Harlan & Zohary 1966). Previous studies have reported that some wild barley accessions maintained in gene bank facilities show evidence of introgression from domesticated barley (Hübner *et al*. 2012; Fang *et al*. 2014; Jakob *et al*. 2014). These studies provide evidence of introgression for individual accessions, but the proportion of the genome introgressed, recency of introgression, and sources of introgression have not been characterized. The geographic ranges of wild and cultivated barley overlap throughout the extensive range of wild barley in the Near East, permitting many opportunities for crop-wild germplasm introgression (Harlan & Zohary 1966). Archeological evidence also demonstrates a long history of human utilization of wild barley as a food source, with evidence of large-scale wild gathering of the grain dating to 23,000 years before present (YBP) (Kislev *et al*. 1992; Weiss *et al*. 2006). Domestication of barley occurred roughly 10,500 YBP (Willcox 2005) over a broad geographic region (Morrell & Clegg 2007) and incorporating multiple wild populations as sources (Poets *et al*. 2015), suggesting an extensive opportunity for both introgression and for populations of wild barley to be altered by the domestication process.

Wild and domesticated barley can be distinguished phenotypically by a relatively small number of canonical domestication and improvement-related traits. These traits are more prevalent in wild barley but are not always exclusive to wild material. Wild barley has brittle spikes at maturity (seed shattering), two rows of fertile florets in the inflorescence, rough barbs on relatively long awns, germination inhibition, and seed dormancy, and generally has more tillers than cultivated barley (Zohary 1969; Pourkheirandish *et al*. 2015). Cultivated barley does not have seed shattering, can be either two-rowed or six-rowed (an improvement trait), has broader leaves and a larger stem diameter, shows no germination inhibition, and can have smoother awns with smaller barbs (Harlan *et al*. 1973; Blumler *et al*. 1991a).

The initial evidence of crop-to-wild introgression often comes from observed phenotypic changes. For example, non-shattering is a characteristic of domesticated barley, but when non-shattering is observed in wild individuals, it suggests introgression has occurred. Accessions selected for Genbank preserved *ex situ* collections typically represent their population of origin in the form of a single accession but carry only a portion of the diversity of the source population and are not suitable for population-level analysis (Schoen & Brown 2001). Most approaches to detect introgression have focused on species and population-level events occurring over many generations (Hibbins & Hahn 2022) and are potentially not applicable here. Thus, our analysis focuses on identifying introgression at the individual accession level.

There are multiple approaches for using molecular markers to detect recent introgression at the individual sample level. The first approach is focused on variants that have large allele frequency differences between a source and recipient population or are private to a source population. For example, Morrell *et al*. (2005) used “cultivar-specific” markers in sorghum to identify weedy johnsongrass individuals showing evidence of introgression. Genetic distance measures relative to labeled populations can also identify a history of hybridization (Pritchard *et al*. 2000). Genetic assignment approaches can also identify admixed individuals subject to introgression using more subtle allele frequency differences among populations. When used with learning samples of individuals with pre-identified labels, genetic assignment approaches are particularly powerful for identifying samples with ancestry derived from one or more populations (Pritchard *et al*. 2000). A potentially more powerful approach to identifying shared ancestry uses genomic segments shared among populations (Ralph & Coop 2013). This approach has previously been applied to barley breeding populations to identify genomic regions introgressed during crossing designed to improve disease resistance (Fang *et al*. 2013) and to identify the relationships among breeding programs (Poets *et al*. 2016). Statistical frameworks to identify genomic segments shared among individuals have continued to improve (Aguillon *et al*. 2022; Browning *et al*. 2023).

Gene bank repositories are intended to preserve plant genetic material and facilitate easy access to these resources for research and plant breeding. The presence of cultivated-to-wild introgression may impact the most appropriate use of individual accessions. Therefore, we applied several evolutionary genetic approaches to the 318 wild barley (*Hordeum vulgare* ssp. *spontaneum)* accessions and 2,446 domesticated (*Hordeum vulgare* ssp. *vulgare*) accessions with SNP array and GBS datasets to address the following questions: 1) What proportion of the genome is introgressed from cultivated to wild barley? 2) Where in the genome does introgression occur, and how large are the introgressed regions? 3) Is the size of introgressed segments suggestive of recent introgression? 4) How often are genes contributing to domestication-related traits introgressed into putatively wild plants? and 5) How is introgression likely to impact inferences based on wild barley genotypic data?

## Materials and Methods

### Plant material

We analyzed 318 wild barley (*Hordeum vulgare* ssp. *spontaneum*) accessions and 2,446 domesticated (*Hordeum vulgare* ssp. *vulgare*) accessions that include both landrace and cultivated barley as defined by passport data from the United States Department of Agriculture, National Small Grain Collection (NSGC) (Muñoz-Amatriaín *et al*. 2014). All of the wild accessions are from the wild barley diversity collection (WBDC) (Steffenson *et al*. 2007), and the domesticated accessions are composed of 803 landraces, and 1,643 cultivated accessions (Poets *et al*. 2015) (Table S1). The wild accessions are primarily from collections at the International Center for Agricultural Research in the Dry Areas (ICARDA), located in Syria, while others are sourced from USDA or IPK gene banks. All accessions were self-fertilized for three to five generations to create inbred lines. Phenotypic data for flag leaf width/length, penultimate leaf width/length, days to heading, plant height, awn length, spike length, kernel number, and tiller number were collected from WBDC accessions grown in common garden environments in the greenhouses in St. Paul, MN (see Supplemental File 1).

### SNP array data

We used 318 WBDC accessions with 3,072 Barley OPA (BOPA) SNPs (Close *et al*. 2009) genotyped on the Illumina platform in Fang *et al*. (2014) (Table 1). Out of these 318 WBDC accessions, 25 WBDC accessions were also genotyped on the barley 9K SNP Illumina iSelect platform (Comadran *et al*. 2012) in Nice *et al*. (2016). There are overlapping SNPs in the BOPA and 9K SNPs. Once duplicates are removed, there are a total of 7,812 SNPs that are polymorphic in both the BOPA and 9K SNP datasets. SNP genotypes for the WBDC were filtered to include polymorphic sites only. SNP quality control procedures for the WBDC accessions followed those in Fang *et al*. (2014) and include removing SNPs with > 15% missing data and > 10% observed heterozygosity based on the rationale that too much missing data could be from inaccurate genotypes and SNPs with high observed heterozygosity are likely errors.

We also utilized 2,446 NSGC Core accessions with 5,010 barley 9K SNPs genotyped in Poets *et al*. (2015). NSGC Core genotyping was filtered to include only polymorphic SNPs. SNPs with > 10% missing data and > 10% observed heterozygosity were removed, similar to criteria in Poets *et al*. (2015). All BOPA and 9K SNP physical positions for the WBDC and NSGC Core accessions were updated relative to Morex_v3 of the reference genome (Mascher *et al*. 2021) by using BLAST comparisons of contextual sequences around each SNP following the methods described in Lei *et al*. (2019) (scripts for updating the physical positions are available in Github at https://github.com/MorrellLAB/morex_reference). For the BOPA and 9K SNPs, we used the consensus genetic maps for the WBDC and NSGC accessions from Muñoz-Amatriaín *et al*. (2011) and (Muñoz-Amatriaín *et al*. 2014).

### Phasing and imputing genotypes

Local ancestry inference analyses (used to test for introgression) required phased variants and no missing genotypes. Thus, we inferred the phase of heterozygous sites and imputed missing genotypes using Beagle v5.4 (Browning *et al*. 2018; Browning *et al*. 2021) for array SNPs that had genetic map positions and physical positions relative to Morex_v3. Beagle uses effective population size as a parameter to determine the hidden Markov Model (HMM) state transition probabilities and recommends specifying appropriate effective population sizes in inbred populations.

Here, wild and domesticated accessions were phased and imputed separately because accessions in this study have been inbred, and there are differences in levels of diversity between wild and domesticated accessions (Morrell *et al*. 2014) and, thus, effective population sizes (*N*_e_). We estimated *N*_e_ of wild and landrace barley from θ = 4*N*_e_μ, where μ is the mutation rate. Here, we use a mutation rate of 5 × 10^-9^ per site (Gaut & Clegg 1993), and θ_W_ (Watterson 1975) estimates of θ_W_ = 5.6 × 10^-3^ in landraces (Morrell *et al*. 2014) and θ_W_ = 8 × 10^-3^ in wild barley (Morrell *et al*. 2006). For phasing and imputation, we used an estimated *N*_e_ = 280,000 for domesticated accessions and *N*_e_ = 400,000 for wild accessions. The parameters for wild and domesticated sets include burn-in = 3 and iterations = 12. All other parameters used Beagle’s defaults. Duplicate markers (where multiple markers had the same cM and physical positions) and markers missing cM locations were excluded. Observed heterozygosity in the datasets was compared before and after phasing and imputation to ensure the heterozygosity level was not inflated due to the imputation process.

### Genotyping-by-sequencing (GBS) data

We also used genotyping-by-sequencing (GBS) data reported by Sallam *et al*. (2017) that includes 314 of the 318 WBDC accessions. GBS data of the WBDC accessions has higher SNP density than the BOPA and 9K genotyping datasets, and because the GBS procedure involves no discovery phase, it can decrease ascertainment bias. DNA extraction, library construction, pooling, sequencing, read alignment, and SNP calling are reported in Sallam *et al*. (2017). GBS variant positions were updated to the Morex_v3 reference genome (Mascher *et al*. 2021). SNPs were filtered to retain only bi-allelic variants. SNPs were removed if GQ < 20, per sample DP < 3 or DP > 70, allele balance deviation was greater than ±10% from 0.5, > 10% heterozygous calls, and > 20% missing genotype calls.

### Inference of ancestral state

We inferred the ancestral state for each SNP in the genotyping datasets using whole-genome resequencing data from *Hordeum murinum* ssp. *glaucum* (Kono *et al*. 2019). We used *H. murinum* ssp. *glaucum* as the outgroup because phylogenetic analyses have placed this diploid species in a clade close to *H. vulgare* (Jakob *et al*. 2004). Raw sequence read processing steps and methods followed those described in Kono *et al*. (2019). After standard sequence read quality control steps and adapter trimming, read alignment was performed using Stampy v1.0.32 (Lunter & Goodson 2011) using prior divergence estimates of 3% with reads mapped against the Morex v3 reference genome (Mascher *et al*. 2021). We generated cleaned BAM files using Samtools v1.9 (Li *et al*. 2009) and Picard v2.20.2 (http://broadinstitute.github.io/picard), then realigned around indels using GATK v3.8.1 (DePristo *et al*. 2011; Van der Auwera & O’Connor 2020). The realigned BAM file was then converted to FASTA format using Analysis of Next-Generation Sequencing Data (ANGSD) v0.931 (Korneliussen *et al*. 2014). Inference of ancestral state was performed using the respective downstream tools used for analyses. For the processing steps above, the following steps were performed using “sequence_handling” (Liu *et al*. 2022) for quality control, adapter trimming, cleaning of BAM files, and realignment around indels. All steps for processing and ancestral state inference are available on GitHub (https://github.com/ChaochihL/Barley_Outgroups).

### Identifying introgressed individuals

Multiple WBDC accessions grown in the greenhouses at the University of Minnesota showed evidence of introgression based on traits characteristic of domesticated barley (Fang *et al*. 2014; Poets *et al*. 2015; Nice *et al*. 2016). This includes WBDC150, WBDC182, WBDC190, and WBDC344, which showed evidence of introgression based on genotyping or the presence of domestication-related traits, such as non-shattering or smooth awns. Multiple accessions from west of the Caspian Sea were excluded from prior studies of wild barley and inference of the origins of domesticated barley because of physical similarities to cultivated barley, such as seed size and erect tillers (Fang *et al*. 2014; Poets *et al*. 2015). This included WBDC150, WBDC205, WBDC217, WBDC227, WBDC228, WBDC229, WBDC231, and WBDC232. We identified additional potentially introgressed accessions based on genetic distances observed in unrooted neighbor-joining trees. We used the PHYLIP v3.695 program “dnadist” to generate distance matrices and then “neighbor” for clustering (Felsenstein 2013). For “dnadist,” we used the F84 model, which incorporates different transition and transversion rates and an input transition/transversion (Ts/Tv) of 1.89 as estimated from the genotype data. Unrooted neighbor-joining (NJ) trees were then visualized with FigTree v1.4.4 (http://tree.bio.ed.ac.uk/software/figtree/). All wild individuals were genotyped with markers developed based on a discovery panel composed primarily of cultivated barley (Close *et al*. 2009; Fang *et al*. 2013), which causes ascertainment bias (Clark *et al*. 2005). This causes greater polymorphism in cultivated accessions that are more similar to the discovery panel (Moragues *et al*. 2010). Thus, wild barley accessions with introgression from cultivated lines are polymorphic for more markers, resulting in longer branch lengths in the NJ tree (see Nice *et al*. (2016)). We also calculated genetic distances using the WBDC GBS dataset. Because variants in the GBS genotyping are identified through *de novo* discovery, there is minimal ascertainment bias relative to the BOPA/9K markers. These distance-based measurements were used to form an initial set of wild accessions with evidence of introgression that could be used in downstream analysis.

To identify additional wild accessions with putative introgression, we used hap-ibd (Zhou *et al*. 2020), which employs a seed-and-extend algorithm to detect segments of shared haplotypes. We ran hap-ibd on the phased and imputed genotype data with the input parameters max-gap = 1000 and min-markers = 40. Then, we pulled only comparisons between one domesticated sample and one wild sample. The list of wild samples that had runs of identity-by-state with domesticated samples was included in our initial query (Table S1).

### Local ancestry inference

We identified the local ancestry of introgressed regions in the full set of domesticated and wild barley genotyping data using FLARE (fast local ancestry estimation) (Browning *et al*. 2023). FLARE was designed for SNP-array and whole genome sequence data, achieves high accuracy compared to existing tools, and scales well to large datasets. FLARE allows for an arbitrary number of ancestries and unknown ancestry relationships among individuals in the reference panel. Phased and imputed genotype data were used for the reference and query panels. Phasing and genotype imputation were performed separately for the wild accessions and domesticated accessions because they were genotyped on different platforms and to avoid artificially inflating heterozygosity in the populations. The domesticated and wild genotype data were combined to form the reference panel, where all SNPs with missing genotypes were removed. Only SNPs with cM positions were included in analyses, and duplicate markers (i.e., the cM and physical positions of SNPs with different IDs were identical) were excluded. For the reference panel, we used the population labels based on passport data: wild, breeding, cultivar, genetic, landrace, and uncertain. Likely introgressed individuals in our query population had the label “wild_introgressed.” We ran FLARE with a random seed under their SNP array data mode. Output from FLARE was then visualized using chromPlot (Oróstica & Verdugo 2016) and ancestry proportions along with other summaries were calculated using a custom R script (available at https://github.com/MorrellLAB/WildIntrogression). Individuals that showed <1% introgressed SNPs were considered “wild” and recycled back into the reference panel wild population. The 1% threshold was chosen from the exploration of the introgressed regions. Individuals with <1% of introgressed SNPs showed introgressed tract lengths that were, on average, 0.4% of the shortest chromosome or 0.05% of the 4.2 Gbp barley genome. When the longest introgressed segment in an individual makes up a small proportion of the genome, it is much less likely from a recent introgression event. We repeated this process until we reached a set of wild individuals that showed >1% of SNPs that had shared ancestry with domesticated individuals.

Individuals with evidence of introgression were treated as a population to test for introgression using the Patterson’s D statistical test (i.e., ABBA-BABA test) (Green *et al*. 2010b; Durand *et al*. 2011). We used the genomics toolset from https://github.com/simonhmartin/genomics_general to run the ABBA-BABA test. We used wild samples for P1, “wild_introgressed” samples for P2, and domesticated samples for P3, and a *Hordeum murinum* sample (Kono *et al*. 2019) as the outgroup (Figure S2). We then tested for gene flow between P3 and P2 (i.e., an excess of ABBA sites relative to BABA) at the genome-wide and per chromosome scales.

### Identifying known domestication-related genes

A literature search was performed to identify known barley genes that are involved in domestication-related traits, such as the non-brittle rachis genes *Btr1* and *Btr2,* which play roles in seed dispersal (Pourkheirandish *et al*. 2015). Other domestication-related traits that differ between wild and domesticated barley include awn and spikelet traits and seed dormancy. We compiled information regarding the GenBank ID, gene function, gene role, and the reporting study (Table S2). Physical locations of the compiled genes were identified via a BLAST alignment against the barley Morex v3 reference genome (Mascher *et al*. 2021) using default values. BLAST results were then filtered by > 95% identity, e-value < 0.0001, and bit score > 60, and the top hit was selected based on the highest percent identity, lowest e-value, and highest bit score. Final hits were processed and reformatted into a standard BED file format. When GenBank IDs did not exist, physical locations of markers identified on the 50k Illumina Infinium iSelect genotyping array (Bayer *et al*. 2017) were used. Only a few 50k markers were identified in published papers and associated with barley genes. For the remaining genes, the trait association between marker haplotypes and loci was used. A haplotype group of adjacent markers can be identified if phenotypic data are available (Xu *et al*. 2023), this approach works for simply inherited morphological traits that can be scored visually. Physical locations of the 50k SNPs relative to Morex_v3 of the reference genome (Mascher *et al*. 2021) were identified with the same methods described above for the BOPA and 9K SNPs.

### Predicting geographic location

Wild barley collections, including the WBDC, have been used extensively to make inferences regarding the geographic origins of domesticated barley, a cultigen derived from at least two domestications (Morrell & Clegg 2007; Pourkheirandish *et al*. 2015; Civáň & Brown 2017). Each accession is representative of the genetic composition of the population of origin. Thus we use a deep-neural network-based approach (Battey *et al*. 2020) implemented in the software locator to infer the geographic origin of introgressed samples based on genotyping data.

Inputs for this analysis are the genotyping data from the BOPA SNPs and the geographic coordinates for samples. The geographic coordinates available for the WBDC are generally limited to two decimal places, thus a resolution of ∼1.1 km. Accessions with evidence of introgression were treated as the query sample, and the balance of accessions were used for training The analysis involved 178 iterations. Uncertainty of sample placement and a 95% confidence interval for sample origin were determined based on 100 bootstrap replicates per query sample.

### Data Availability

Genotyping data for the 318 WBDC accessions are available in the Data Repository for the University of Minnesota (DRUM), http://doi.org/10.13020/D6B59N. The raw genotyping data for 2,446 landrace and cultivated accessions are available in FigShare, https://figshare.com/articles/dataset/Raw_Genotyping_Data_Barley_landraces_are_characterized_by_geographically_heterogeneous_genomic_origins/1468432. Additional files that are too include as supplemental tables will be available through DRUM and are available for review at https://drive.google.com/drive/folders/1-79yKHm_TkNgOJjbYZO4QA7TSLlBSS0V?usp=sharing. Scripts for data processing and analysis are available in the GitHub repository: https://github.com/MorrellLAB/WildIntrogression.

## Results

### Identification of introgression from domesticated barley into wild relatives

Previous studies have shown evidence for crop-to-wild introgression in barley (Fang *et al*. 2014; Poets *et al*. 2015; Nice *et al*. 2016). Of the 318 WBDC accessions, 14 wild accessions were initially thought to contain introgressions from domesticated accessions (defined here as either landraces or cultivated accessions) (Table S1). These were accessions identified as showing introgression in previous studies based on allele frequency differences, genetic distance observed in NJ trees, or phenotypic observations when the plants were grown in the greenhouse. Using FLARE, we identified 48 wild individuals with introgression from domesticated accessions (Figure 1), hereafter wild-introgressed accessions. Eight of the 48 wild-introgressed individuals identified by FLARE were previously shown to be introgressed (Table S1), as indicated by wild samples with longer branches in NJ trees based on genetic distance measures (Fang *et al*. 2014; Nice *et al*. 2016). In the GBS dataset of wild individuals, some of the longer NJ tree branches corresponded with wild individuals identified in the genotype dataset, although the difference was less pronounced (Figure S1).

**Figure 1.**
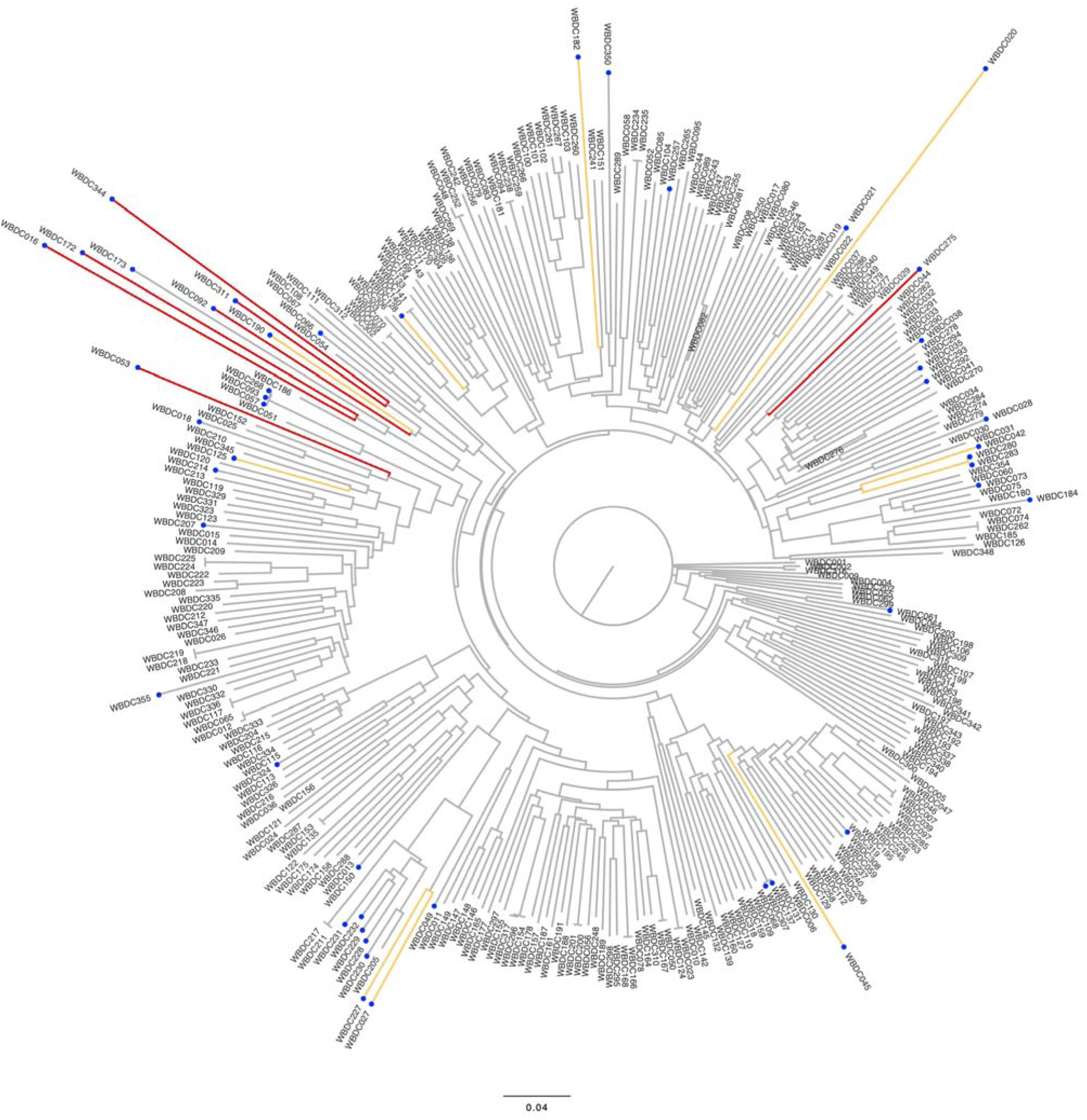
An unrooted neighbor-joining tree of wild genotyped accessions based on the WBDC BOPA dataset. Blue circles are wild-introgressed samples identified by FLARE (local ancestry inference). Yellow indicates accessions with ≥ 20 Mbp tracts of introgression, and red indicates accessions with ≥ 50 Mbp tracts of introgression. The tracts were identified by FLARE using introgressed regions with the full set of domesticated and wild barley SNP genotyping data.

### Characterization of introgression

Local ancestry inference with FLARE (Browning *et al*. 2023) identified an initial set of 48 wild-introgressed lines, where a subset was also identified by the NJ tree (Figure 2). Here, we defined wild-introgressed accessions as individuals with > 1% of SNPs identified as sharing ancestry with a domesticated individual, and they will be used in the following analyses. The rest of the samples were relabeled as “wild.” In the 48 wild-introgressed accessions, there was an average of 58.7 continuous segments (tracts) identified in each sample (Table S3), with a total of 2,817 tracts of introgression (overlapped with 18,149 high-confidence genes). The average size of introgressed tracts genome-wide was 1.7 Mbp ± 8.1, and the average size per chromosome ranged from 0.96-3.3 Mbp (Figure 3; Table S4). There were 11 tracts ≥ 50 Mbp across nine samples, six segments ≥ 100 Mbp across six samples, and one tract ≥ 200 Mbp in one sample. The largest introgressed tract was 224.2 Mbp on chr1H in WBDC190 (Figure 4, the complete set of chromosome plots will be available in a DRUM repository). The introgressed tract ≥ 100 Mbp were identified in the accessions WBDC016, WBDC092, WBDC182, WBDC190, WBDC275, and WBDC344 (for a complete list of accessions and the size of the introgression tracts, see Table S5). Generally, long introgressed tracts are thought to have occurred recently because there should be fewer opportunities for recombination to break up tracts of introgression (Martin & Jiggins 2017). Occasionally, some longer tracts can occur by chance due to the stochastic nature of inheritance (Ralph & Coop 2013). Still, tracts > 50 Mbp are relatively rare (0.4% of all tracts identified) and likely not due to historical shared ancestry.

**Figure 2.**
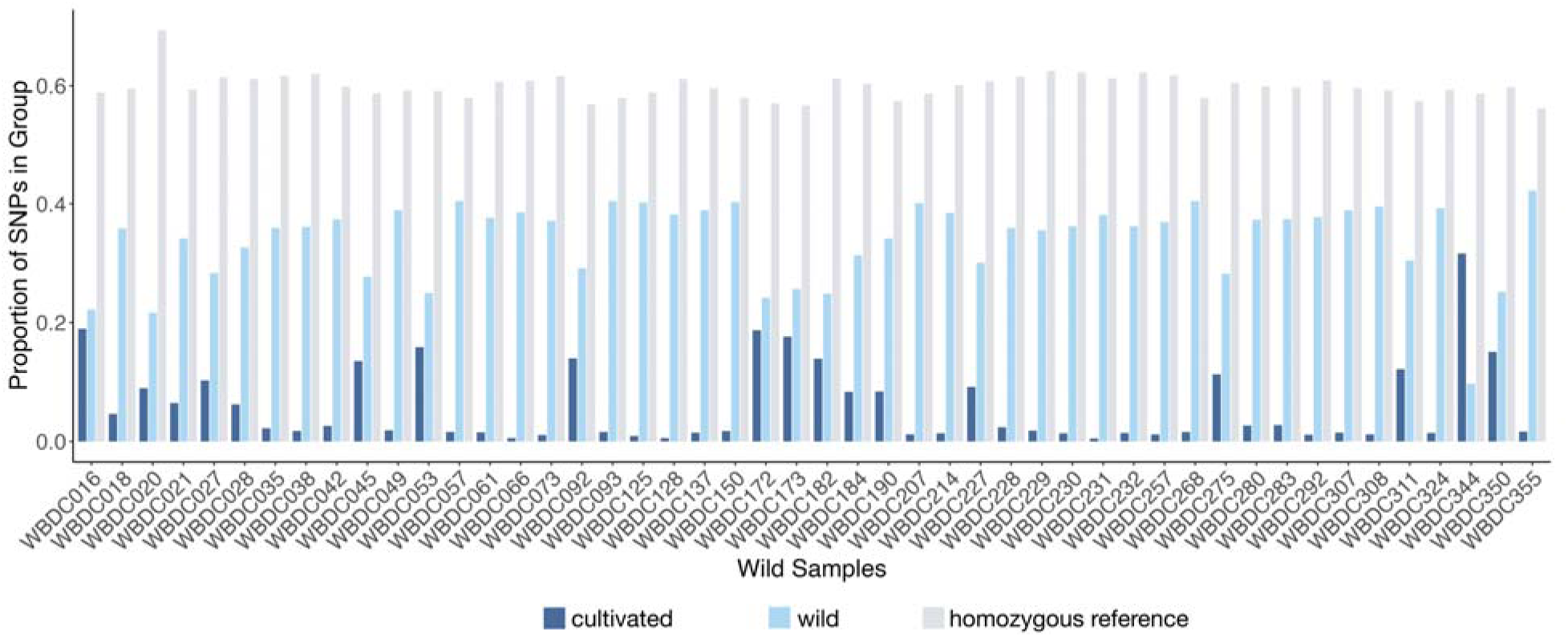
The proportion of SNPs in wild accessions that were assigned domesticated ancestry. Light blue and dark blue represent alternate genotypes assigned to either wild or cultivated (categorized as domesticated) populations. Grey represents homozygous reference genotypes that were not counted towards introgressed segments.

**Figure 3.**
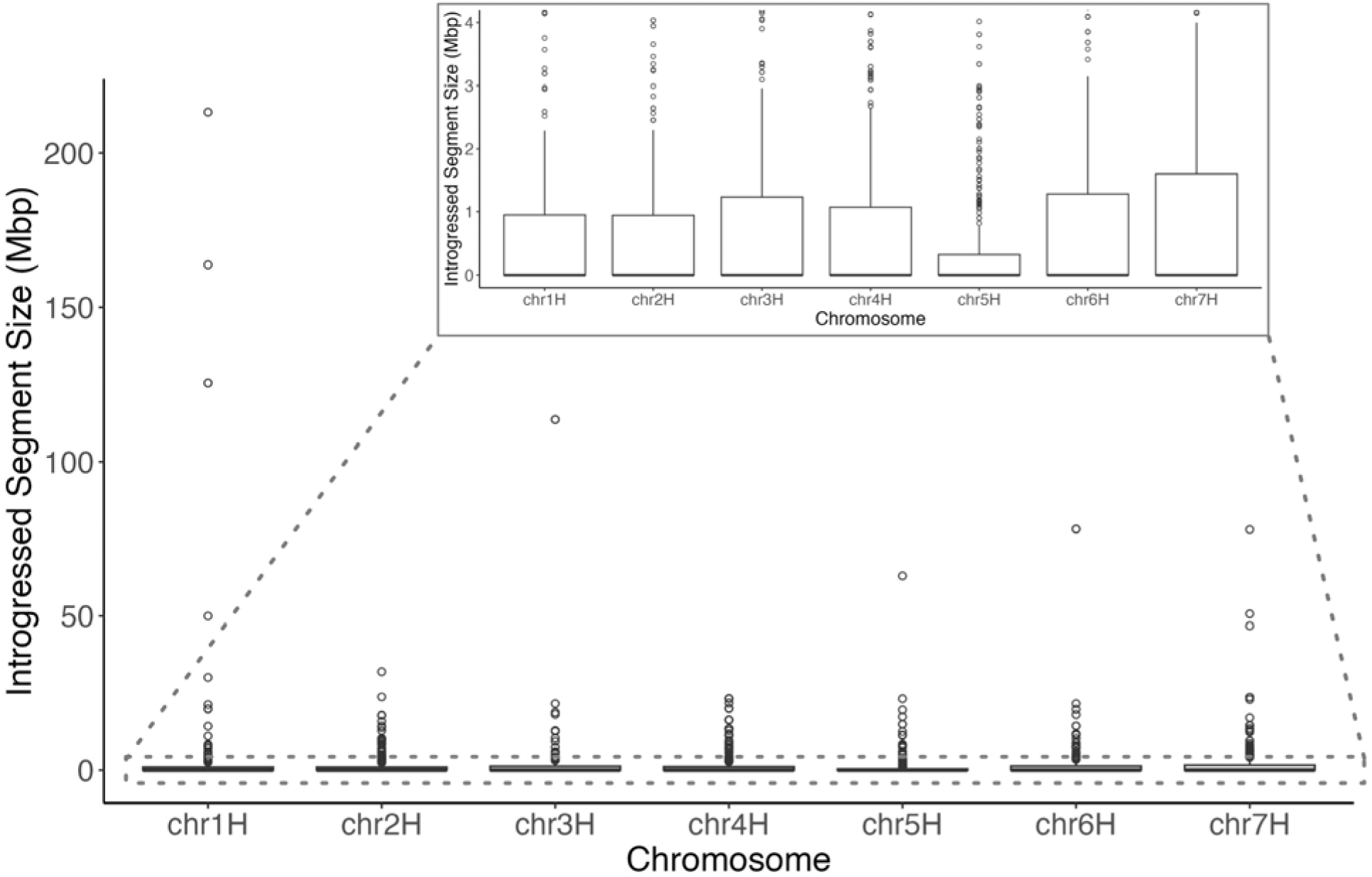
The distributions of domesticated to wild introgressed segment lengths on each chromosome for all samples. The inset depicts the first 4 Mbp of segments.

**Figure 4.**
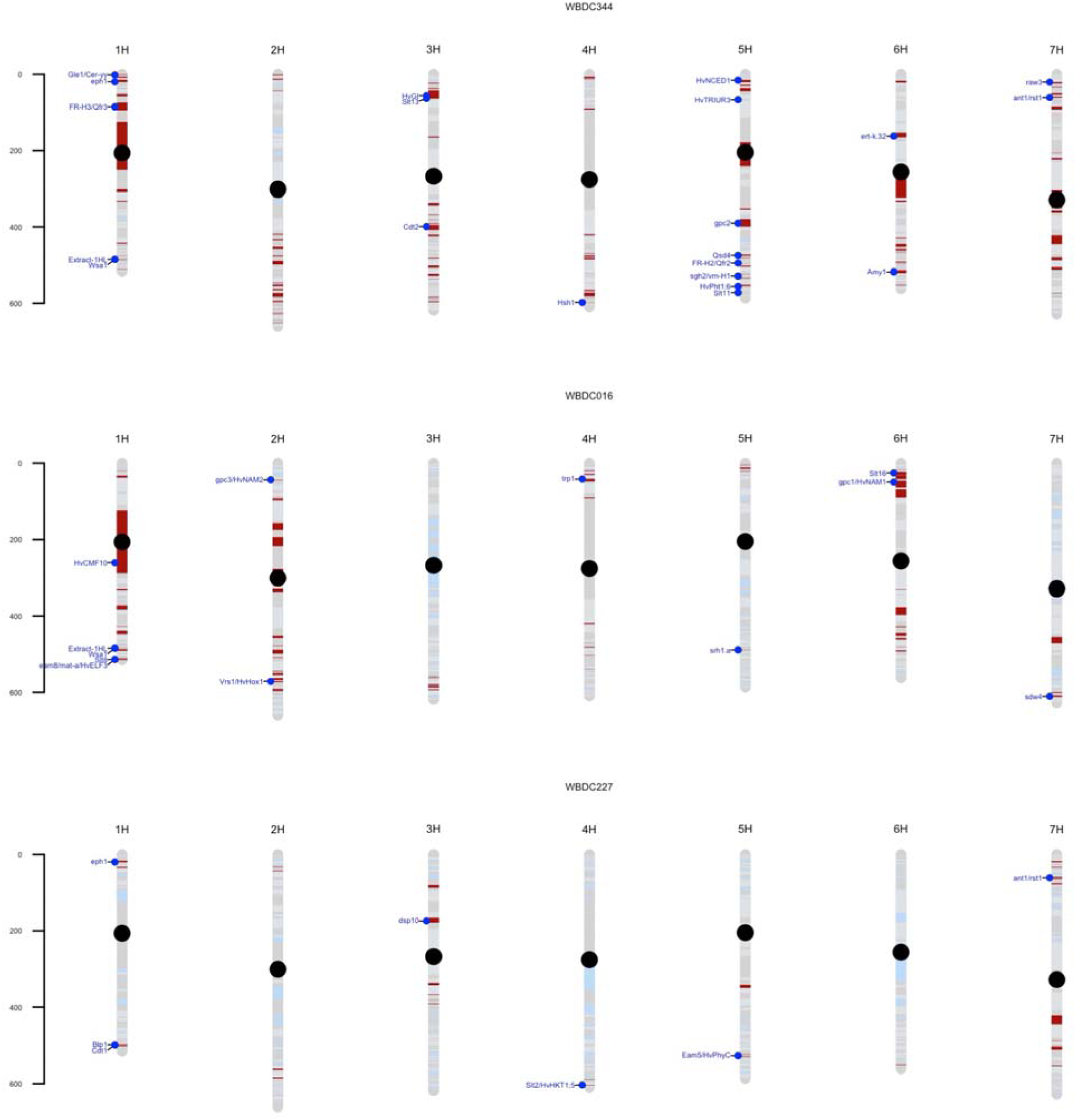
Introgressed segments are shown for three representative samples. Red segments are introgressed regions from domesticated accessions, and light blue segments have wild ancestry. The dot represents centromeres for each chromosome. Annotations are gene name abbreviations (Table S2) when domestication-related genes overlap introgressed regions.

For wild-introgressed accessions, we assessed evidence of gene flow from domesticated accessions into wild accessions using Patterson’s D statistic, the ABBA-BABA test (Green *et al*. 2010a). Our ABBA-BABA test resulted in a genome-wide excess of ABBA sites over BABA (*D* = 0.083, Z = 5.12) when P1 = wild, P2 = wild_introgressed, P3 = domesticated, and P4 = outgroup (Figure S2). This suggests more shared derived alleles between domesticated samples (P3) and wild-introgressed samples (P2), indicating gene flow between the populations. Based on chromosome-level ABBA tests, all chromosomes except for chr5H and chr6H show evidence of gene flow between wild-introgressed and domesticated samples (Table S6).

### Introgression and known domestication-related genes

To identify loci where introgression from cultivated barley could impact wild barley populations, we used a literature search to identify 184 barley genes associated with a variety of phenotypic changes (Table S2). We focus on domestication-related genes, based on the definitions of Harlan *et al*. (1973). We do not include disease resistance related genes in our primary comparisons because it is often not clear from published reports, including mutant screens, if there is naturally occurring variation that differentiates wild and cultivated populations. This resulted in a subset of 125 genes included in downstream analyses, which will be referred to as “domestication-related” genes from here on. Of the cloned domestication-related genes identified, 20 genes (16% of domestication-related genes) overlapped introgressed segments (Table S5). In wild-introgressed accessions, we identified 25 individuals where at least one introgressed tract overlapped a domestication-related gene (Figure S3). WBDC344 had the most introgressed tracts that overlap with domestication-related genes, some of which include seed dormancy-related genes (HvNCED1, Qsd4) (Figure 4). WBDC344 was an individual initially visually identified as wild-introgressed with the characteristics non-shattering, smooth awn, Intermedium-C (modifier of VRS1, which regulates two-row versus six-row formation) (Ramsay *et al*. 2011), and looked semi-dwarf. The overlap between introgressed segments and domestication-related loci do not directly explain the observed phenotypes. This may be partly due to the multigenic nature of many traits; only segments of the genome carry domestication and improvement-related alleles, while the bulk of ancestry across the genome derives from wild ancestry. Also, many of the currently cloned and well-studied genes are involved in crop improvement rather than the early stages of domestication (see Table S2 for descriptions of gene functions). The three other individuals (WBDC150, WBDC182, WBDC190) visually identified as showing evidence of introgression were all identified as having introgressed segments based on identity-by-state/identity-by-descent, local ancestry inference, and NJ tree approaches.

### Wild introgressed barley phenotypes

To determine if there are phenotypes that can help distinguish between wild individuals and wild-introgressed individuals, we looked at phenotype data collected on the WBDC accessions (Supplemental File 1). Wider leaves are characteristic of domesticated barley (Figure 5). Flag leaf width in introgressed individuals appears slightly wider than in wild lines but is not significantly different (*P*-value = 0.06485, Wilcoxon rank sum test). Awn lengths appear slightly longer in the wild compared to introgressed individuals, but again, these differences were not significant (*P*-value = 0.5635, Wilcoxon rank sum test). For other measured traits, including flag leaf length, penultimate leaf length, plant height, days to heading, kernel number, and tiller number, there were no discernible differences between wild and introgressed individuals (Figure S4).

**Figure 5.**
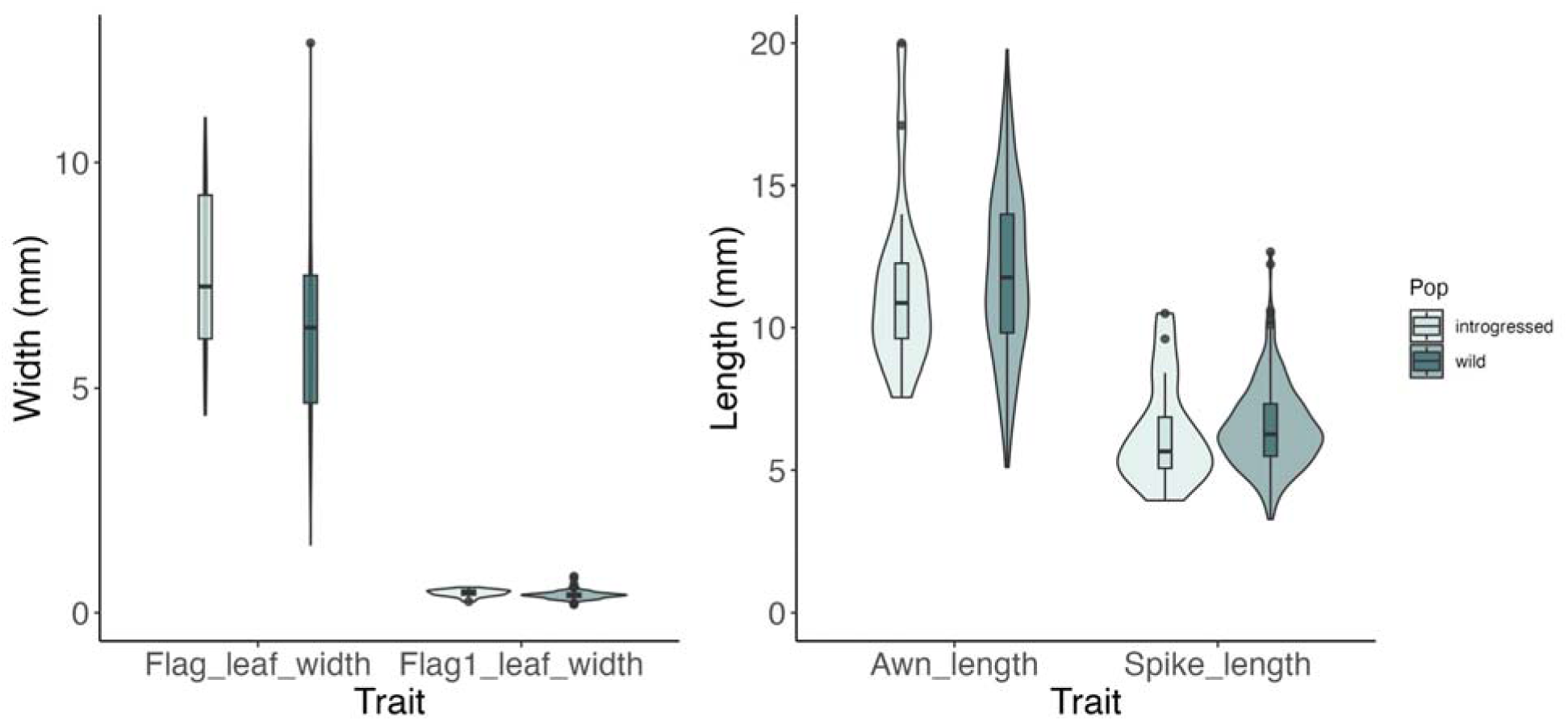
The distributions of the traits flag leaf width, penultimate leaf width (Flag1_leaf_width), awn length, and spike length in wild-introgressed versus wild barley accessions. Wild introgressed accessions here are those with ≥ 21 Mbp introgressed tract lengths on average. Domesticated barley typically have wider leaves and shorter awn lengths than wild barley.

### Impact of introgression on genetic composition

We used genotyping data and estimation of location of origin for introgressed samples to determine how genetic composition was altered by introgression. Accessions with the highest proportion of introgression demonstrated relatively large geographic displacements. WBDC016 and WBDC344 have estimated cultivated introgression in 11.6% and 16.6% of the genome, respectively (Table S3), and both demonstrate relatively large shifts in geographic placement (Figure 6). WBDC016, for example, is shifted from a sample location in the southern Zagros Mountains to a 95% confidence interval of placement in the Southern Levant, closer to the wild populations that are most similar to Western cultivated barley (Poets *et al*. 2015). Alternatively, samples with very low levels of introgression have an inferred 95% confidence interval of origin centered on their collection location; see WBDC061 and WBDC231 in Figure 6. For some samples, notably those from the Caspian Sea region, intermediate levels of introgression result in very broad 95% confidence intervals of geographic placement based on genotypes (Figure S5).

**Figure 6.**
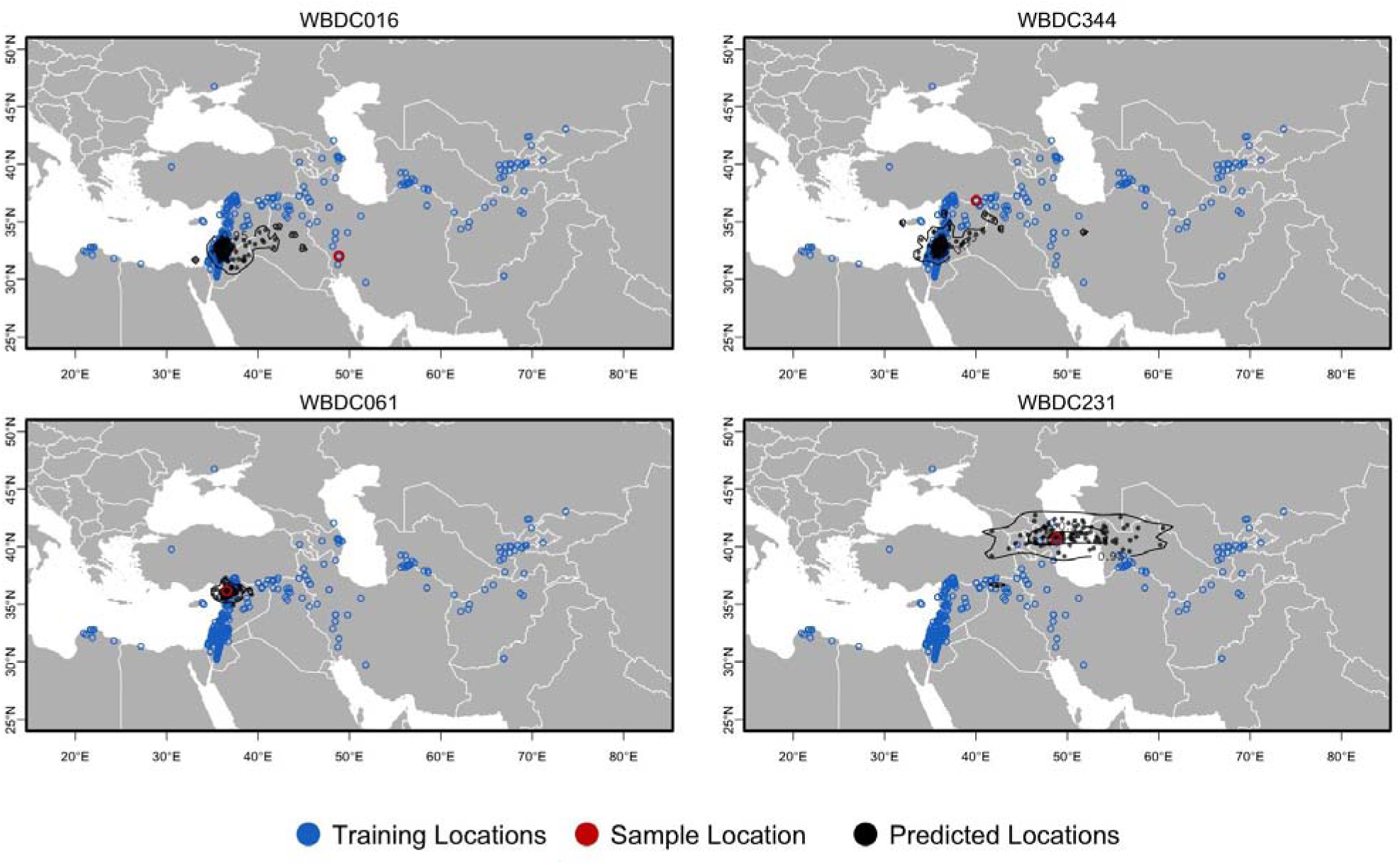
Predicted geographic location of WBDC genomes using the locator tool. The figure depicts the uncertainty in the geographic location predictions for each WBDC sample using 100 bootstrap replicates. Blue points correspond to sample locations in the training set. The red circles indicate the true geographic location of each query sample. Black points are predicted locations generated from bootstrap replicates, and black lines outline the 95% confidence intervals based on bootstrap replicates.

## Discussion

Introgression between cultivated plants and wild relatives is common (Ellstrand 2003) and can impact the genetic composition of wild populations (Hufford *et al*. 2013; Wang *et al*. 2023), with implications for the conservation of natural populations (Wang *et al*. 2023). Hybridization is particularly likely when cultigens and their wild relatives occur in sympatry, are reproductively intercompatible, and have similar chromosomal composition. Thus it is not surprising that hybrids between wild and cultivated barley are commonly observed in natural populations (Harlan & Zohary 1966). Consistent with previous studies (Hübner *et al*. 2012; Fang *et al*. 2014; Jakob *et al*. 2014; Nice *et al*. 2016), we identified individuals showing evidence of domesticated introgression into wild barley.

On average, 6% ± 6.8% of SNPs in each wild introgressed individual were inferred to be derived from domesticated samples, which make up on average 2.4% ± 3.3% of the 4.2 Gbp genome (Mascher *et al*. 2021). The largest total amount of introgression appeared in WBDC344 (16.6% of the genome identified as introgressed) (Figure S2). Although many of the introgressed wild samples identified showed relatively short tracts of introgression, individuals with more obvious domestication-related traits typically have larger tracts of introgression. This is most likely because individuals with larger tracts of introgression tend to have more introgressed segments overall (Figure S6), and thus more opportunities for domestication-related genes to be present.

All four WBDC individuals identified as potentially introgressed based on the presence of cultivated barley phenotypes were confirmed to be subject to introgression from cultivated barley (see Figure 2). Among eight additional accessions from west of the Caspian Sea that resemble cultivated barley in plant architecture, six show evidence of introgression, with only WBDC205 (from southern Russia) and WBDC217 (from Armenia) not appearing to be introgressed. Introgressed samples from this region show moderate levels of introgression (Figure 2).

Local recombination rate impacts the size of regions introgressed in experimental wild to cultivated barley populations, with large tracts introgressed in low recombination regions (Dreissig *et al*. 2020). Larger tracts of introgression tend to occur near the centromere, whereas shorter segments tend to occur towards the distal ends of the chromosomes. This is likely due to the pericentromere showing reduced rates of recombination in barley (Künzel *et al*. 2000; Baker *et al*. 2014; Lei *et al*. 2019). Self-fertilization in wild barley after an initial hybridization event limits the opportunity to break up larger introgressed segments, particularly in genomic regions with a low crossover rate (Chakraborty & Smouse 1988; Martin & Jiggins 2017).

First-generation hybrids contain complete chromosomes from the cultivated donor. Despite selfing rate of >98.2% in both cultivated (Abdel-Ghani *et al*. 2004) and wild barley (Brown *et al*. 1978), effective recombination in early generations after hybridization will break introgressed segments into smaller tracts of introgression. The median size of introgressed tracts is 31.4 Mbp, with a small number of tracts in the 125-225 Mbp size range, relative to barley chromosomes that are >500 Mbp in length (Figure 4). Among the 48 introgressed individuals, the median number of introgressed segments is 21.5 (Table S3). Because the WBDC is an ex-situ collection, there is potential for hybridization to occur during seed replication in gene banks, and there is evidence that introgression is more common in gene bank lines than in wild populations (Jakob *et al*. 2014). It is possible that some of the introgression observed occurred during seed replication. However, chromosome-scale tracing of introgressed tracts is also highly suggestive of introgression in natural environments before seed collection for preservation at the gene bank facilities. Also, georeferenced locality (passport) data places many of the wild barley source populations near grain fields, orchards, or from locations with a high density of irrigated agriculture. Given the proximity between wild and cultivated barley since initial domestication, it is plausible that much of the introgression observed involves retaining introgressed segments from hybridization in natural populations.

### Introgressed regions and domestication genes

We found 50% of the high-confidence genes annotated in the Morex v3 reference genome overlapped introgressed regions in at least one wild barley line. We identified a set of 125 genes related to domestication or improvement in barley. Of these, 38 were cloned genes with a reference sequence interval that could be defined based on nucleotide sequence positions. Introgressed wild barley lines can carry many domestication-related alleles (Figure 4) while showing minimal trait-based evidence of introgression. This is likely due to the multigenic nature of many traits, such that introgression from cultivated sources remains cryptic because only a portion of genes that modify a trait are from domesticated sources.

### The lack of diagnostic traits for introgression

We find phenotypic differences between wild-introgressed barley and wild barley, but no single trait or combination of traits is diagnostic of introgression (Figure 5; Figure S3). Wild barley has extensive phenotypic diversity (Steffenson *et al*. 2007), with desert ecotypes (Volis *et al*. 2002) for example, that are phenotypically distinct from plants from other regions. One trait that could diagnose introgression is a non-shattering phenotype (characteristic of domesticates) observed in a wild individual (Harlan *et al*. 1973; Blumler *et al*. 1991b). However, the presence of non-shattering is insufficient to identify most individuals with significant introgression.

### Wild introgressed samples as a genetic resource

The wild-introgressed individuals remain valuable genetic resources because a large proportion of the genome of each accession carries wild alleles. Lines with relatively small tracts of introgression can be treated as near-isogenic lines, which isolate genomic segments from cultivated barley in a wild barley genetic background.

### Introgression and evolutionary inference hybridization

Previous studies have emphasized the role of hybridization of wild and cultivated barley during the maintenance of ex-situ collections (Jakob *et al*. 2014) and the potential for recent introgression to impact studies of evolutionary history and genetic provenance of domesticates. Given that wild and cultivated barley are grown in close proximity during seed production in gene banks, hybridization is quite likely. However, our results identify many relatively small genomic regions impacted by introgression (Figure 4), consistent with older introgression events preserved in inbreeding lineages.

Though there is extensive migration across the range of wild barley, with haplotypes at many genes shared species-wide, significant population structure is evident at a large proportion of loci (Morrell *et al*. 2003; Morrell & Clegg 2007). The majority of haplotypes at any locus occur in both wild and cultivated barley (Morrell *et al*. 2014; Schmid *et al*. 2018), but introgression from cultivated lines has the potential to introduce haplotypes that differ from those that predominate in the recent, coalescent history of local populations. For samples with the largest degree of introgression, including WBDC016 and WBDC344, this will shift the genetic composition sufficiently to dramatically change the genotype-based inference of location of origin. WBDC016 has an inferred location of origin ∼800 km west of the collection site, while WBDC344 is shifted ∼650 km to the southwest (Figure 6). Among samples with low levels of introgression, the effect on geographic placement is modest (Figure 6; Figure S5).

The effects of including introgressed samples in comparative analyses will vary based on the nature of the analysis. For many applications, such as inferring the geographic origins of domestication-related traits by identifying extant wild accessions that carry haplotypes that are the most similar to those associated with domestication or improvement allele (Komatsuda *et al*. 2007; Pourkheirandish *et al*. 2015), introgression will cause greater variance in the apparent geographic distribution of haplotypes among wild populations. WBDC355 (also known as OUH602), has often been used as a comparator in studies cloning individual barley genes (Nair *et al*. 2010). OUH602 also has published genome assembly(Sato *et al*. 2021). We found 0.198% introgression from cultivated lines, but this did not result in the presence of domesticated barley genomic segments at any of the loci examined here (Figure S7).

The introduction of haplotypes through introgression could also create admixture linkage disequilibrium (Chakraborty & Smouse 1988; Briscoe *et al*. 1994) and extended haplotype homozygosity (Voight *et al*. 2006). Extensive introgression, such that introduced haplotypes predominate, could result in false positive signals of selection. Nucleotide sequence diversity and population differentiation, as measured by F_ST_, could also be altered by introgression, with the introduction of new haplotypes increasing the variance in the distribution of coalescence times and thus also inference based on the number of segregating sites or pairwise diversity in the sample (Austerlitz *et al*. 1997). Introgression will also reduce the degree of allele frequency differentiation among populations, potentially impacting studies seeking to identify adaptive variants using environmental association. One study seeking to identify adaptive variants using environmental association and F_ST_ outliers Fang *et al*. (2014) reported excluding 14 of the 318 WBDC accessions where introgression was most evident. Poets *et al*. (2015) reported culling the WBDC to 284 accessions to avoid including individuals with introgression in a panel used to infer the geographic contributions of wild barley populations to cultivated barley. Relatively aggressive removal of introgressed samples is likely warranted in studies focused on wild populations expected to have a long history of local adaptation. Also, for applications such as identifying targets of selection, it would be preferable to exclude wild introgressed individuals.

There is a long history of using advanced backcross populations to incorporate segments of the wild barley genome into cultivated lines (Pillen *et al*. 2003; Maurer *et al*. 2015; Nice *et al*. 2017). These studies typically report the introgression of wild barley into cultivated lines, followed by backcrossing to cultivated lines that are more tractable as a study population and have potentially desirable agronomic traits. Samples with some degree of introgression may not be particularly disruptive in genetic mapping studies where mixing of wild and cultivated ancestry is a primary goal (Pillen *et al*. 2003; Maurer *et al*. 2015; Nice *et al*. 2017).

Identification of introgressed line can inform decisions on plant material preservation in gene banks. This could include renewed efforts at collecting wild lines in regions from which most current accessions are introgressed. Our study found that introgression can go undetected or is only detectable through genetic means, but understanding the levels of introgression present in populations can help researchers make informed decisions related to genetic resource preservation.

## Acknowledgments

We thank Nadia Janis for compiling the initial list of domestication-related loci and Malik Samuels for growing and maintaining the barley plants in the greenhouse. Elaine Lee provided assistance with updating figures. This study was supported by a University of Minnesota Informatics Institute MnDRIVE Graduate Assistantship award to Chaochih Liu, the National Science Foundation (grant IOS-1339393), and the Minnesota Agricultural Experiment Station fund (MIN-13-122 in support of Peter L. Morrell). This research was carried out with software and hardware support provided by the Minnesota Supercomputing Institute (MSI) at the University of Minnesota.

## Author Contributions

CL and PLM wrote the paper. LL performed the early genotypic data integration and analysis, idea development, and revision of the manuscript. CL and JBP performed the analyses. MS, JBP, and JDF compiled the list of domestication-related loci and genomic intervals. BJS, JCS, JKR, AHS, and RTG collected the phenotypic data on the WBDC samples. JDF and PLM visually identified wild plants that showed traits suggesting introgression.

## Supplemental Figures

**Figure S1.**
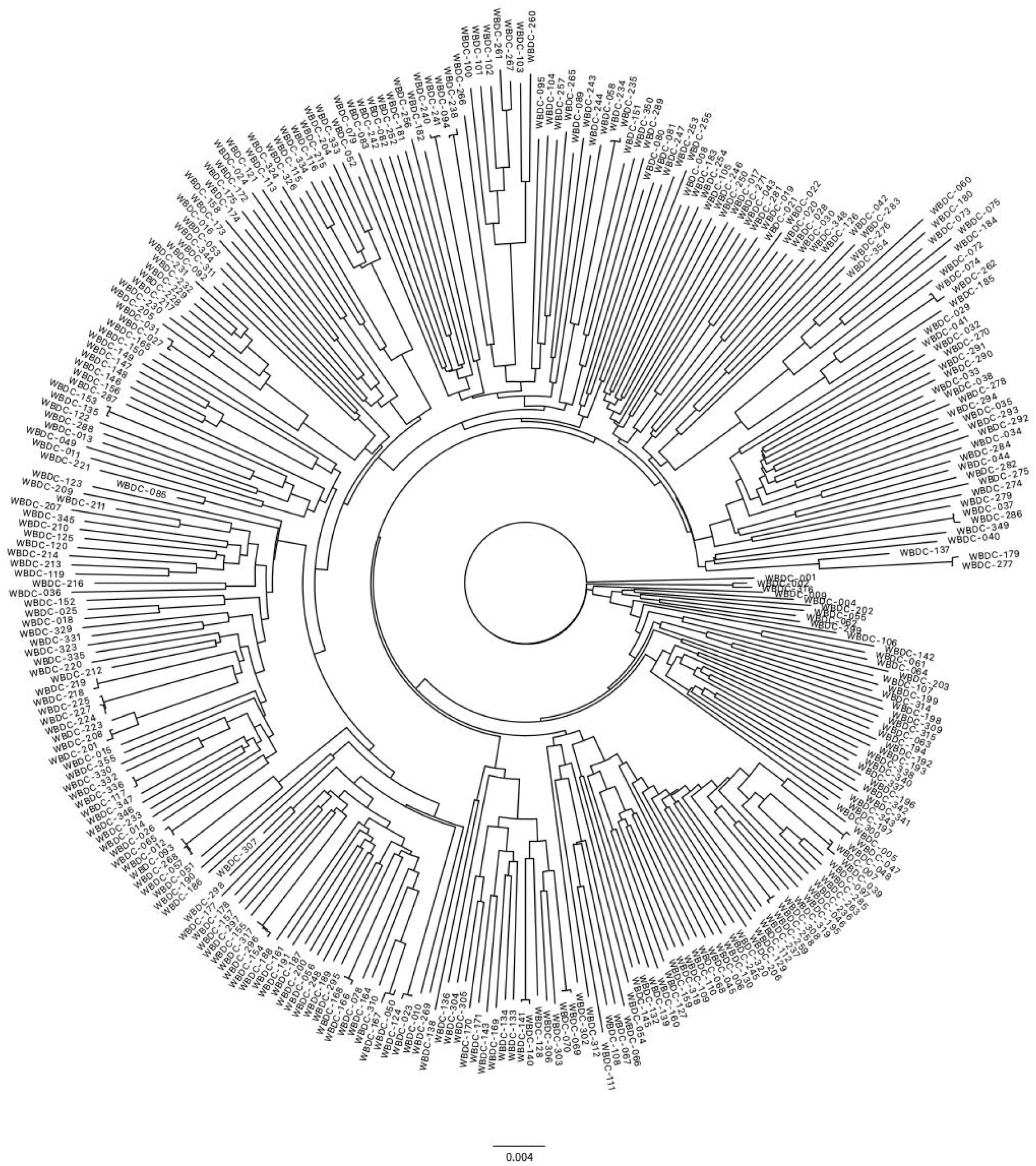
Unrooted neighbor-joining tree of wild genotyped accessions based on genotyping-by-sequencing.

**Figure S2.**
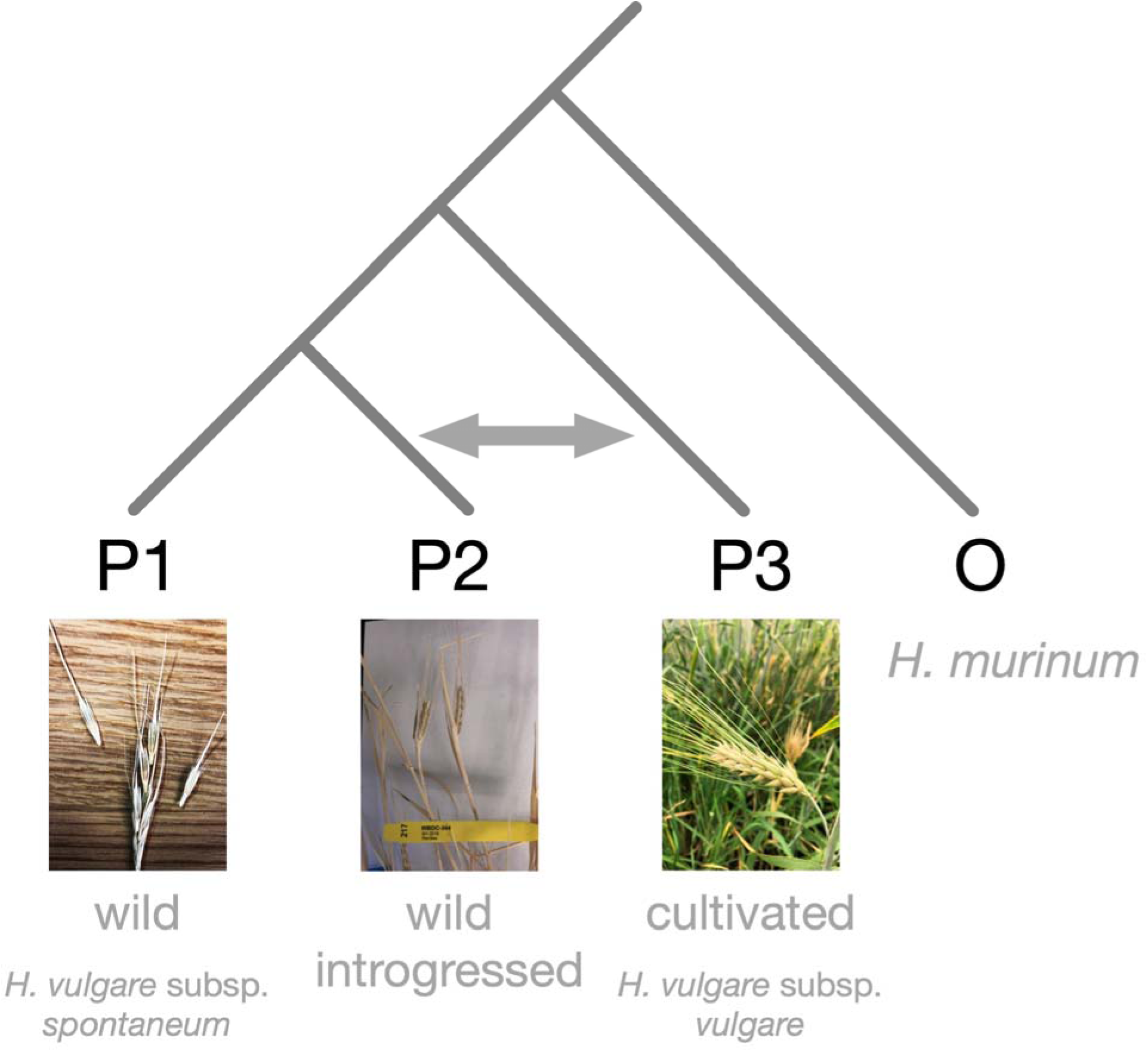
Photos depicting characteristic phenotypic differences between wild and domesticated barley. The cartoon depicts an ABBA-BABA test for gene flow, treating wild introgressed individuals as a population.

**Figure S3.**
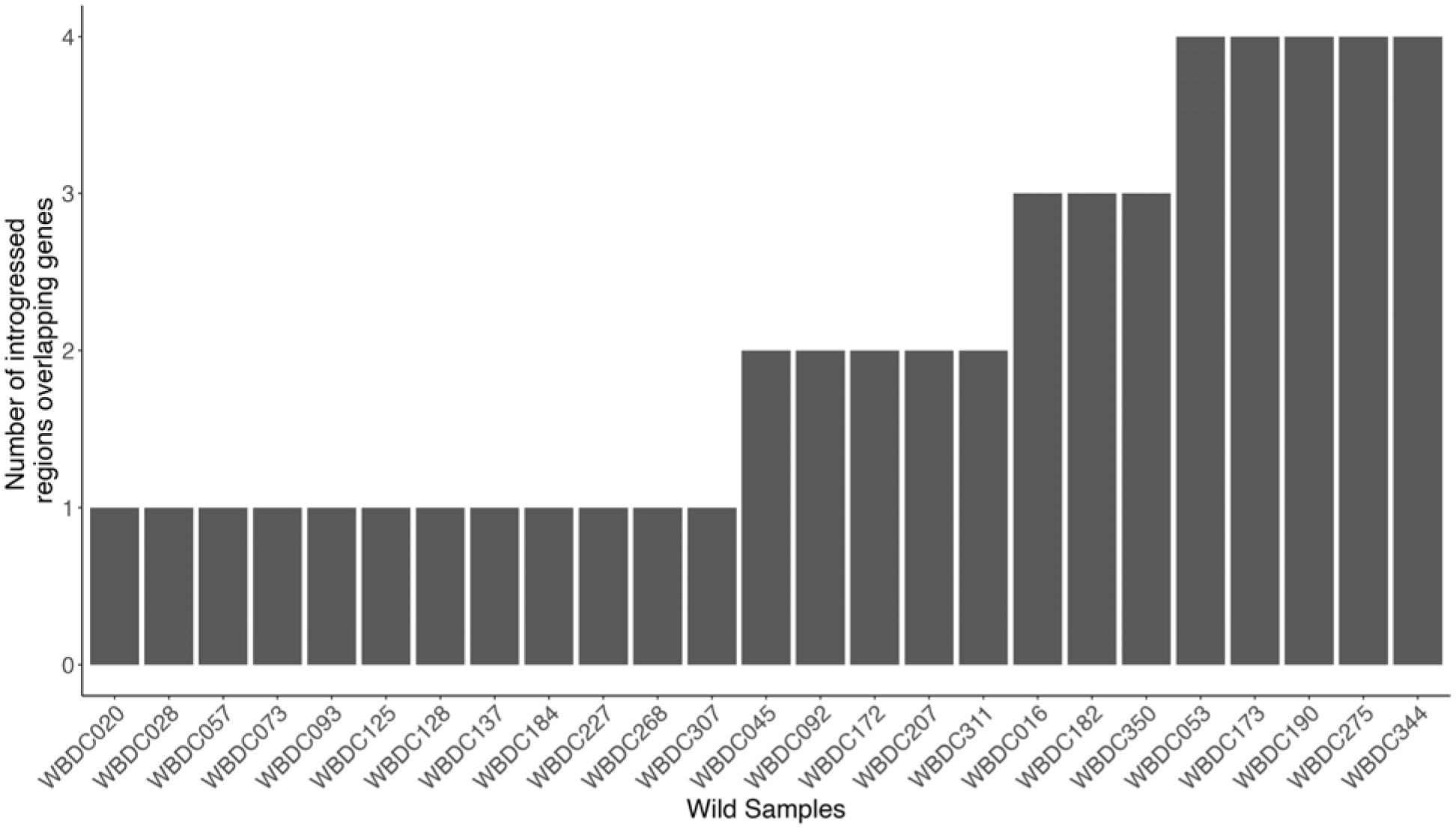
The number of introgressed segments in wild individuals that overlap cloned barley genes potentially involved in domestication.

**Figure S4.**
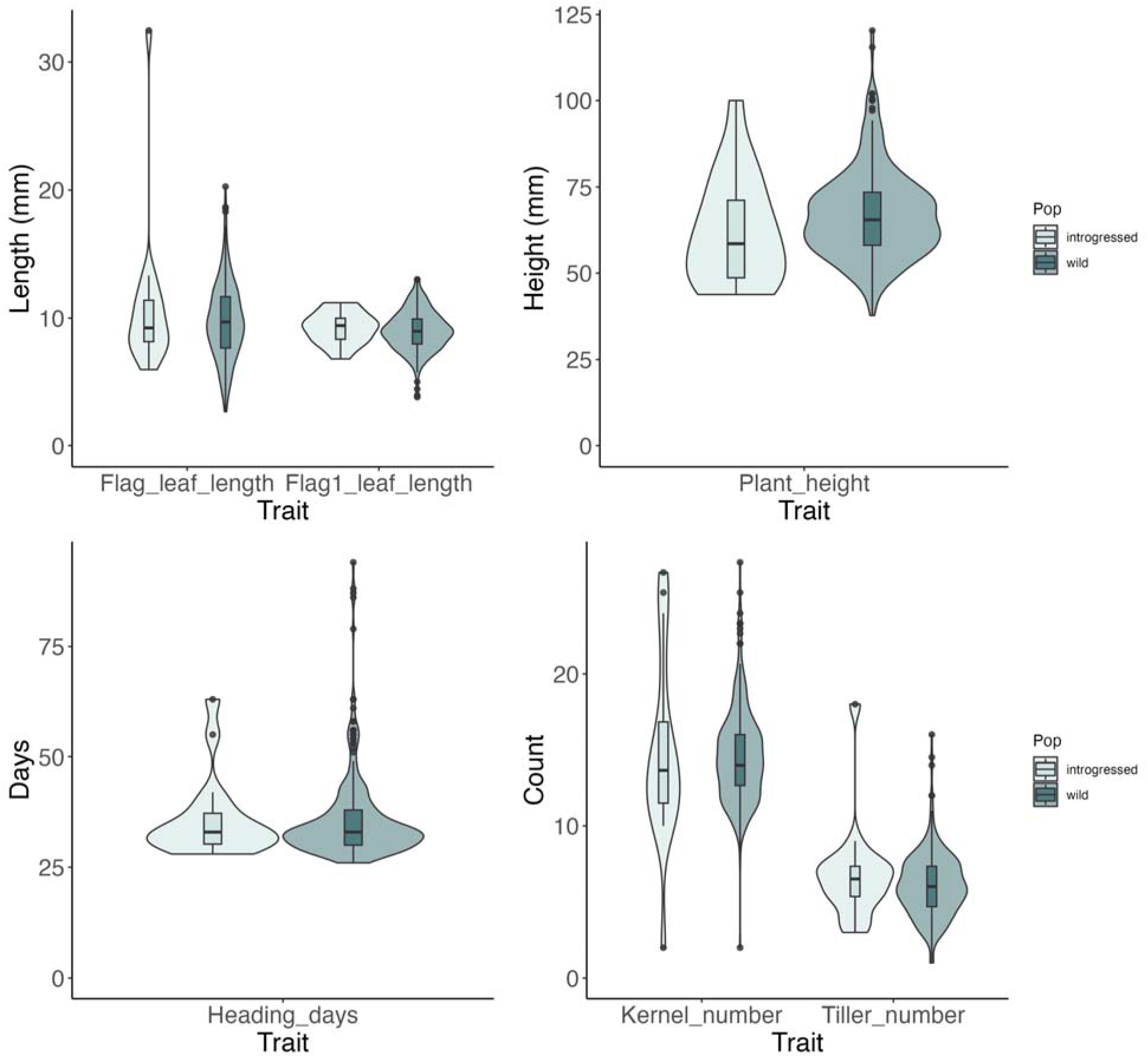
Distributions of the traits flag leaf length, penultimate leaf length (Flag1_leaf_length), plant height, days to heading, kernel number, and tiller number in wild-introgressed versus wild barley accessions. Wild introgressed accessions here are those with ≥ 21 Mbp introgressed tract lengths on average.

**Figure S5.**
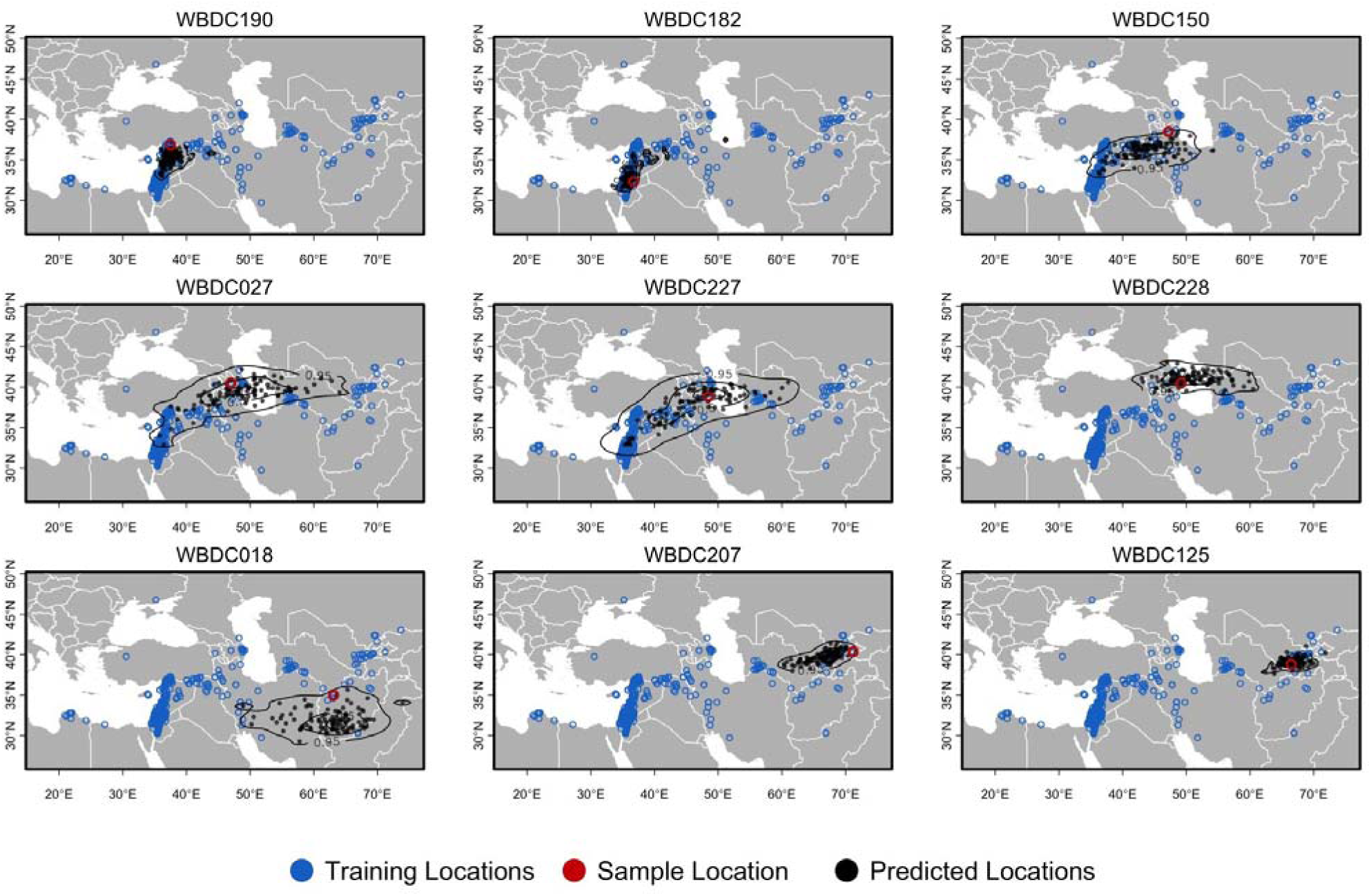
Predicted geographic location of WBDC genomes using locator tool. Shows uncertainty in the geographic location predictions for each WBDC sample using a bootstrap analysis approach. The predictions are derived from 100 bootstrap samples. Blue point corresponds to a predicted location generated from a bootstrap, red marks true geographic location of sample and black shows region where 95% of bootstrap prediction occurred.

**Figure S6.**
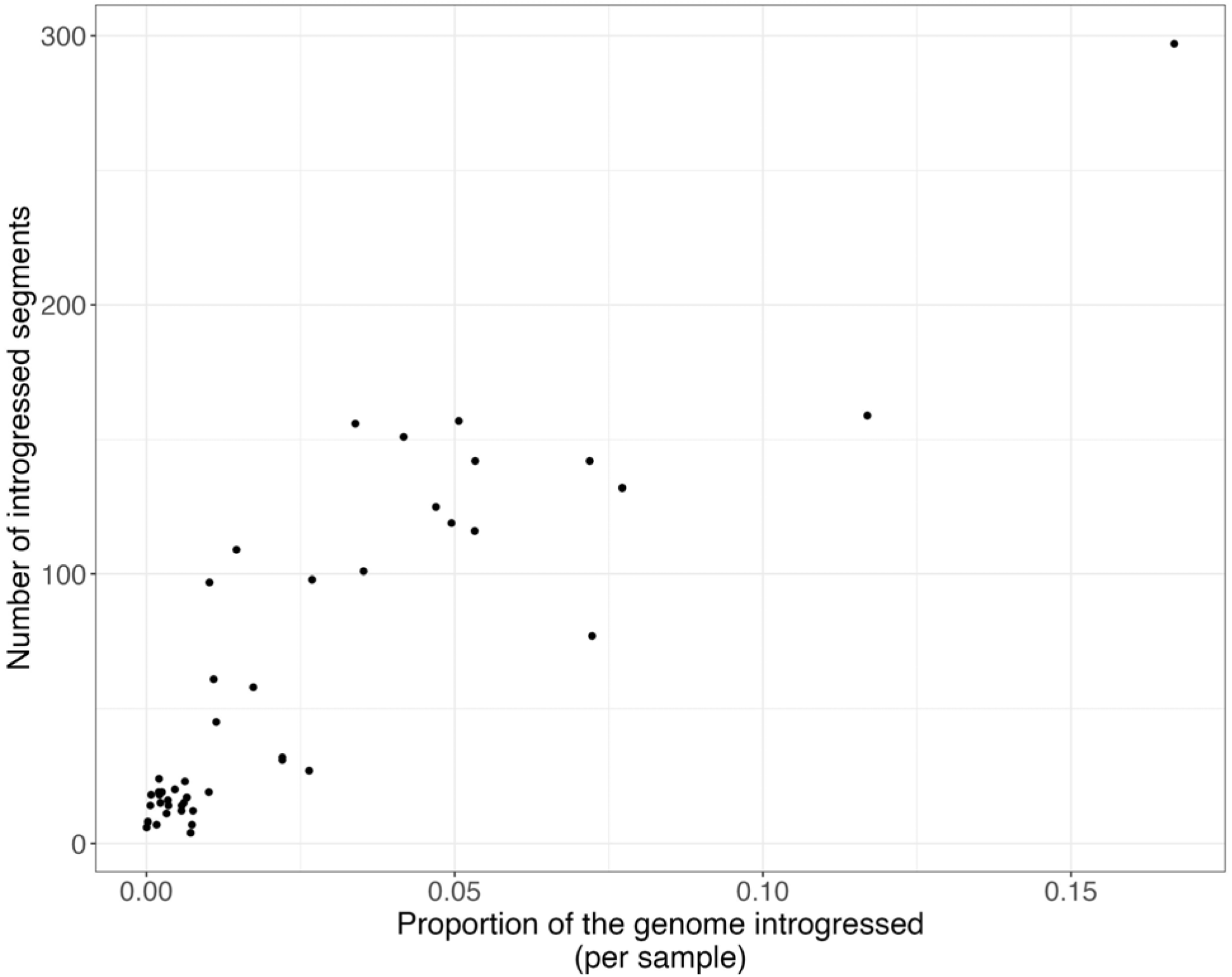
The proportion of the genome identified as introgressed in each wild individual compared to the number of introgressed segments identified in each sample.

**Figure S7.**
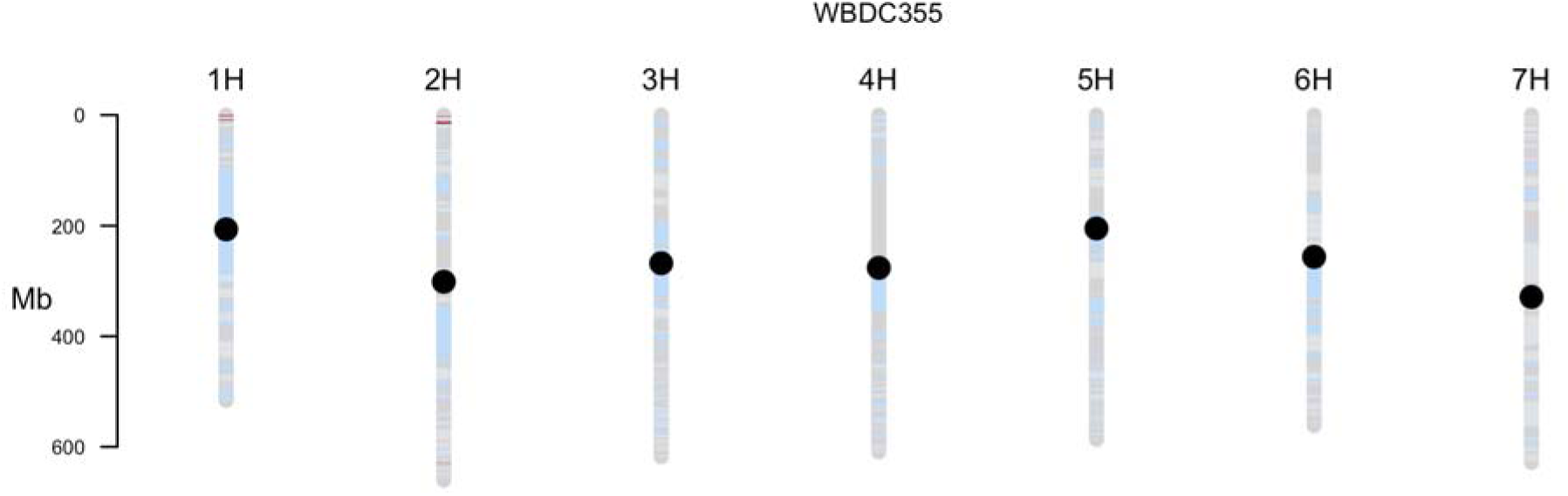
Introgressed segments are shown for WBDC355 (OUH602). Red segments are introgressed regions from domesticated accessions, and light blue segments have wild ancestry. The dot represents centromeres for each chromosome. Annotations are gene name abbreviations (Table S2) when domestication-related genes overlap introgressed regions.

## Notes

https://drive.google.com/drive/folders/1-79yKHm_TkNgOJjbYZO4QA7TSLlBSS0V?usp=sharing

### Competing Interest Statement

The authors have declared no competing interest.

## References

Abdel-Ghani AH, Parzies HK, Omary A, Geiger HH (2004) Estimating the outcrossing rate of barley landraces and wild barley populations collected from ecologically different regions of Jordan. Theoretical and Applied Genetics, 109, 588–595.

Aguillon SM, Dodge TO, Preising GA, Schumer M (2022) Introgression. Current Biology, 32, R865–R868.

Austerlitz F, Jung-Muller B, Godelle… B (1997) Evolution of coalescence times, genetic diversity and structure during colonization. Theoretical population …,

Baker K, Bayer M, Cook N et al. (2014) The low-recombining pericentromeric region of barley restricts gene diversity and evolution but not gene expression. The Plant Journal, 79, 981–992.

Battey CJ, Ralph PL, Kern AD (2020) Predicting geographic location from genetic variation with deep neural networks. Elife, 9, e54507.

Bayer MM, Rapazote-Flores P, Ganal M et al. (2017) Development and evaluation of a barley 50k iSelect SNP array. Frontiers in Plant Science, 8, 1792.

Blumler MA, Byrne R, Belfer-Cohen A et al. (1991a) The ecological genetics of domestication and the origins of agriculture [and comments and reply]. Current Anthropology, 32, 23–54.

Blumler MA, Byrne R, Belfer-Cohen A et al. (1991b) The ecological genetics of domestication and the origins of agriculture [and comments and reply]. Current Anthropology, 32, 23–54.

Bohra A, Kilian B, Sivasankar S et al. (2022) Reap the crop wild relatives for breeding future crops. Trends in Biotechnology, 40, 412–431.

Briscoe D, Stephens JC, O’Brien SJ (1994) Linkage disequilibrium in admixed populations: applications in gene mapping. Journal of Heredity, 85, 59–63.

Brown AHD, Zohary D, Nevo E (1978) Outcrossing rates and heterozygosity in natural populations of *Hordeum spontaneum* Koch in Israel. Heredity, 41, 49–62.

Browning BL, Tian X, Zhou Y, Browning SR (2021) Fast two-stage phasing of large-scale sequence data. American Journal of Human Genetics, 108, 1880–1890.

Browning BL, Zhou Y, Browning SR (2018) A one-penny imputed genome from next-generation reference panels. American Journal of Human Genetics, 103, 338–348.

Browning SR, Waples RK, Browning BL (2023) Fast, accurate local ancestry inference with FLARE. American Journal of Human Genetics, 110, 326–335.

Chakraborty R, Smouse PE (1988) Recombination of haplotypes leads to biased estimates of admixture proportions in human populations. Proceedings of the National Academy of Sciences of the United States of America, 85, 3071–3074.

Civáň P, Brown TA (2017) A novel mutation conferring the nonbrittle phenotype of cultivated barley. New Phytologist, 214, 468–472.

Clark AG, Hubisz MJ, Bustamante CD, Williamson SH, Nielsen R (2005) Ascertainment bias in studies of human genome-wide polymorphism. Genome Research, 15, 1496–1502.

Close TJ, Bhat PR, Lonardi S et al. (2009) Development and implementation of high-throughput SNP genotyping in barley. BMC Genomics, 10, 1–13.

Comadran J, Kilian B, Russell J et al. (2012) Natural variation in a homolog of Antirrhinum *CENTRORADIALIS* contributed to spring growth habit and environmental adaptation in cultivated barley. Nature Genetics, 44, 1388–1392.

Dempewolf H, Krishnan S, Guarino L (2023) Our shared global responsibility: safeguarding crop diversity for future generations. Proceedings of the National Academy of Sciences of the United States of America, 120, e2205768119.

DePristo MA, Banks E, Poplin R et al. (2011) A framework for variation discovery and genotyping using next-generation DNA sequencing data. Nature Genetics, 43, 491.

Dreissig S, Maurer A, Sharma R et al. (2020) Natural variation in meiotic recombination rate shapes introgression patterns in intraspecific hybrids between wild and domesticated barley. New Phytologist, 228, 1852–1863.

Durand EY, Patterson N, Reich D, Slatkin M (2011) Testing for ancient admixture between closely related populations. Molecular Biology and Evolution, 28, 2239–2252.

Ellstrand NC (2003) Dangerous liaisons? When cultivated plants mate with their wild relatives.

Ellstrand NC, Meirmans P, Rong J et al. (2013) Introgression of crop alleles into wild or weedy populations. Annual Review of Ecology, Evolution, and Systematics, 44, 325–345.

Fang Z, Eule-Nashoba A, Powers C et al. (2013) Comparative analyses identify the contributions of exotic donors to disease resistance in a barley experimental population. G3: Genes, Genomes, Genetics, 3, 1945–1953.

Fang Z, Gonzales AM, Clegg MT et al. (2014) Two genomic regions contribute disproportionately to geographic differentiation in wild barley. G3: Genes, Genomes, Genetics, 4, 1193–1203.

Felsenstein J (2013) PHYLIP (Phylogeny Inference Package) version 3.695. Distributed by the author,

Flint-Garcia S, Feldmann MJ, Dempewolf H, Morrell PL, Ross-Ibarra J (2023) Diamonds in the not-so-rough: Wild relative diversity hidden in crop genomes. PLoS biology, 21, e3002235.

Gaut BS, Clegg MT (1993) Molecular evolution of the *Adh1* locus in the genus *Zea*. Proceedings of the National Academy of Sciences of the United States of America, 90, 5095–5099.

Green RE, Krause J, Briggs AW et al. (2010a) A draft sequence of the Neandertal genome. Science, 328, 710–722.

Green RE, Krause J, Briggs AW et al. (2010b) A draft sequence of the Neandertal genome. Science, 328, 710–722.

Harlan JR, de Wet JMJ, Price EG (1973) Comparative evolution of cereals. Evolution, 27, 311–325.

Harlan JR, Zohary D (1966) Distribution of wild wheats and barley. Science, 153, 1074–1080.

Hibbins MS, Hahn MW (2022) Phylogenomic approaches to detecting and characterizing introgression. Genetics, 220, iyab173.

Hübner S, Günther T, Flavell A et al. (2012) Islands and streams: clusters and gene flow in wild barley populations from the Levant. Molecular Ecology, 21, 1115–1129.

Hufford MB, Lubinksy P, Pyhäjärvi T, Devengenzo MT, Ellstrand NC, Ross-Ibarra J (2013) The genomic signature of crop-wild introgression in maize. PLoS Genet, 9, e1003477.

Jakob SS, Rödder D, Engler JO et al. (2014) Evolutionary history of wild barley (*Hordeum vulgare* subsp. *spontaneum*) analyzed using multilocus sequence data and paleodistribution modeling. Genome Biology and Evolution, 6, 685–702.

Jakob SS, Meister A, Blattner FR (2004) The considerable genome size variation of *Hordeum* species (Poaceae) is linked to phylogeny, life form, ecology, and speciation rates. Molecular Biology and Evolution, 21, 860–869.

Kislev ME, Nadel D, Carmi I (1992) Epipalaeolithic (19,000 BP) cereal and fruit diet at Ohalo II, Sea of Galilee, Israel. Review of Palaeobotany and Palynology, 73, 161–166.

Komatsuda T, Pourkheirandish M, He C et al. (2007) Six-rowed barley originated from a mutation in a homeodomain-leucine zipper I-class homeobox gene. Proceedings of the National Academy of Sciences, 104, 1424–1429.

Kono TJY, Liu C, Vonderharr EE et al. (2019) The fate of deleterious variants in a barley genomic prediction population. Genetics, 213, 1531–1544.

Korneliussen TS, Albrechtsen A, Nielsen R (2014) ANGSD: analysis of next generation sequencing data. BMC Bioinformatics, 15, 356.

Künzel G, Korzun L, Meister A (2000) Cytologically integrated physical restriction fragment length polymorphism maps for the barley genome based on translocation breakpoints. Genetics, 154, 397–412.

Lei L, Poets AM, Liu C et al. (2019) Environmental association identifies candidates for tolerance to low temperature and drought. G3: Genes, Genomes, Genetics, 9, 3423–3438.

Li H, Handsaker B, Wysoker A et al. (2009) The sequence alignment/map format and SAMtools. Bioinformatics, 25, 2078–2079.

Liu C, Hoffman P, Wyant S et al. (2022) MorrellLAB/sequence_handling: Release v3.0: SNP calling with GATK 4.1 and Slurm compatibility (v3.0.0).

Lunter G, Goodson M (2011) Stampy: a statistical algorithm for sensitive and fast mapping of Illumina sequence reads. Genome Research, 21, 936–939.

Martin SH, Jiggins CD (2017) Interpreting the genomic landscape of introgression. Current Opinion in Genetics & Development, 47, 69–74.

Mascher M, Wicker T, Jenkins J et al. (2021) Long-read sequence assembly: a technical evaluation in barley. Plant Cell, 33, 1888–1906.

Maurer A, Draba V, Jiang Y, Schnaithmann… F (2015) Modelling the genetic architecture of flowering time control in barley through nested association mapping. BMC Genomics,

McCouch SR, Rieseberg LH (2023) Harnessing crop diversity. Proceedings of the National Academy of Sciences of the United States of America, 120, e2221410120.

Moragues M, Comadran J, Waugh R, Milne I, Flavell AJ, Russell JR (2010) Effects of ascertainment bias and marker number on estimations of barley diversity from high-throughput SNP genotype data. Theoretical and Applied Genetics, 120, 1525–1534.

Morrell PL, Gonzales AM, Meyer KKT, Clegg MT (2014) Resequencing data indicate a modest effect of domestication on diversity in barley: a cultigen with multiple origins. Journal of Heredity, 105, 253–264.

Morrell PL, Lundy KE, Clegg MT (2003) Distinct geographic patterns of genetic diversity are maintained in wild barley (Hordeum vulgare ssp. spontaneum) despite migration. Proceedings of the National Academy of Sciences, 100, 10812–10817.

Morrell PL, Toleno DM, Lundy KE, Clegg MT (2006) Estimating the contribution of mutation, recombination and gene conversion in the generation of haplotypic diversity. Genetics, 173, 1705–1723.

Morrell PL, Clegg MT (2007) Genetic evidence for a second domestication of barley (*Hordeum vulgare*) east of the Fertile Crescent. Proceedings of the National Academy of Sciences of the United States of America, 104, 3289–3294.

Morrell PL, Williams-Coplin TD, Lattu AL, Bowers JE, Chandler JM, Paterson AH (2005) Crop-to-weed introgression has impacted allelic composition of johnsongrass populations with and without recent exposure to cultivated sorghum. Molecular Ecology, 14, 2143–2154.

Muñoz-Amatriaín M, Cuesta-Marcos A, Endelman JB et al. (2014) The USDA barley core collection: genetic diversity, population structure, and potential for genome-wide association studies. PloS One, 9, e94688.

Muñoz-Amatriaín M, Moscou MJ, Bhat PR, et al. (2011) An improved consensus linkage map of barley based on flow-sorted chromosomes and single nucleotide polymorphism markers. The Plant Genome, 4,

Nair SK, Wang N, Turuspekov Y et al. (2010) Cleistogamous flowering in barley arises from the suppression of microRNA-guided HvAP2 mRNA cleavage. Proceedings of the National Academy of Sciences, 107, 490–495.

Nice LM, Steffenson BJ, Blake TK, Horsley RD, Smith KP, Muehlbauer GJ (2017) Mapping agronomic traits in a wild barley advanced backcross–nested association mapping population. Crop Science, 57, 1199–1210.

Nice LM, Steffenson BJ, Brown-Guedira GL et al. (2016) Development and genetic characterization of an advanced backcross–nested association mapping (AB-NAM) population of wild × cultivated barley. Genetics, 203, 1453–1467.

Oróstica KY, Verdugo RA (2016) chromPlot: visualization of genomic data in chromosomal context. Bioinformatics, 32, 2366–2368.

Pillen K, Zacharias A, Léon J (2003) Advanced backcross QTL analysis in barley (Hordeum vulgare L.). Theoretical and Applied Genetics, 107, 340–352.

Poets AM, Mohammadi M, Seth K et al. (2016) The effects of both recent and long-term selection and genetic drift are readily evident in North American barley breeding populations. G3: Genes, Genomes, Genetics, 6, 609–622.

Poets AM, Fang Z, Clegg MT, Morrell PL (2015) Barley landraces are characterized by geographically heterogeneous genomic origins. Genome Biology, 16, 173.

Pourkheirandish M, Hensel G, Kilian B et al. (2015) Evolution of the grain dispersal system in barley. Cell, 162, 527–539.

Pritchard JK, Stephens M, Donnelly P (2000) Inference of population structure using multilocus genotype data. Genetics, 155, 945–959.

Ralph P, Coop G (2013) The geography of recent genetic ancestry across Europe. PLoS Biol, 11, e1001555.

Ramsay L, Comadran J, Druka A et al. (2011) *INTERMEDIUM-C*, a modifier of lateral spikelet fertility in barley, is an ortholog of the maize domestication gene *TEOSINTE BRANCHED 1*. Nature Genetics, 43, 169–172.

Rieseberg LH, Wendel JF (1993) Introgression and its consequences in plants. Hybrid zones and the evolutionary process, 70, 109.

Sallam AH, Tyagi P, Brown-Guedira G, Muehlbauer GJ, Hulse A, Steffenson BJ (2017) Genome-wide association mapping of stem rust resistance in *Hordeum vulgare* subsp. *spontaneum*. G3: Genes, Genomes, Genetics, 7, 3491–3507.

Sato K, Mascher M, Himmelbach A, Haberer G, Spannagl M, Stein N (2021) Chromosome-scale assembly of wild barley accession “OUH602”. G3, 11, jkab244.

Schmid K, Kilian B, Russell J (2018) Barley domestication, adaptation and population genomics. The barley genome, 317–336.

Schoen DJ, Brown AHD (2001) The conservation of wild plant species in seed banks: attention to both taxonomic coverage and population biology will improve the role of seed banks as conservation tools. Bioscience, 51, 960–966.

Steffenson BJ, Olivera P, Roy JK, Jin Y, Smith KP, Muehlbauer GJ (2007) A walk on the wild side: mining wild wheat and barley collections for rust resistance genes. Australian Journal of Agricultural Research, 58, 532–544.

Van der Auwera GA, O’Connor BD (2020) Genomics in the cloud: using Docker, GATK, and WDL in Terra.

Voight BF, Kudaravalli S, Wen X, Pritchard JK (2006) A map of recent positive selection in the human genome. PLoS biology,

Volis S, Mendlinger S, Ward D (2002) Differentiation in populations of *Hordeum spontaneum* Koch along a gradient of environmental productivity and predictability: plasticity in response to water and nutrient stress. Biological Journal of the Linnean Society, 75, 301–312.

Wang N, Cao S, Liu Z et al. (2023) Genomic conservation of crop wild relatives: A case study of citrus. PLoS genetics, 19, e1010811.

Watterson GA (1975) On the number of segregating sites in genetical models without recombination. Theoretical Population Biology, 7, 256–276.

Weiss E, Kislev ME, Hartmann A (2006) Autonomous cultivation before domestication. Science, 312, 1608–1610.

Willcox G (2005) The distribution, natural habitats and availability of wild cereals in relation to their domestication in the Near East: multiple events, multiple centres. Vegetation History and Archaeobotany, 14, 534–541.

Xu T, Meng S, Zhu X et al. (2023) Integrated GWAS and transcriptomic analysis reveal the candidate salt-responding genes regulating Na+/K+ balance in barley (Hordeum vulgare L.). Frontiers in Plant Science, 13, 1004477.

Zhou Y, Browning SR, Browning BL (2020) A fast and simple method for detecting identity-by-descent segments in large-scale data. The American Journal of Human Genetics, 106, 426–437.

Zohary D (1969) The progenitors of wheat and barley in relation to domestication and agricultural dispersal in the Old World. (eds Ucko PJ, Dimbleby GW), pp. 47–66. Duckworth and Company, London.

